# LncRNA-dependent nuclear stress bodies promote intron retention through SR protein phosphorylation

**DOI:** 10.1101/681056

**Authors:** Kensuke Ninomiya, Shungo Adachi, Tohru Natsume, Junichi Iwakiri, Goro Terai, Kiyoshi Asai, Tetsuro Hirose

## Abstract

A number of long noncoding RNAs (lncRNAs) are induced in response to specific stresses to construct membrane-less nuclear bodies; however, their function remains poorly understood. Here, we report the role of nuclear stress bodies (nSBs) formed on highly repetitive satellite III (HSATIII) lncRNAs derived from primate-specific satellite III repeats upon thermal stress exposure. A transcriptomic analysis revealed that depletion of HSATIII lncRNAs, resulting in elimination of nSBs, promoted splicing of 533 retained introns during thermal stress recovery. A HSATIII-Comprehensive identification of RNA-binding proteins by mass spectrometry (ChIRP-MS) analysis identified multiple splicing factors in nSBs, including serine and arginine-rich pre-mRNA splicing factors (SRSFs), the phosphorylation states of which affect splicing patterns. SRSFs are rapidly dephosphorylated upon thermal stress exposure. During stress recovery, CDC like kinase 1 (CLK1) was recruited to nSBs and accelerated the re-phosphorylation of SRSF9, thereby promoting target intron retention. Our findings suggest that HSATIII-dependent nSBs serve as a conditional platform for phosphorylation of SRSFs by CLK1 to promote the rapid adaptation of gene expression through intron retention following thermal stress exposure.

## Introduction

Long noncoding RNAs (lncRNAs) have recently been recognized as fundamental regulators of gene expression, but their mechanisms of action remain largely unknown. Among the tens of thousands of human lncRNAs, a subset of architectural RNAs (arcRNAs) function as structural scaffolds of membrane-less subnuclear organelles or nuclear bodies (NBs) (Chujo, Yamazaki et al., 2016). NBs are usually located in inter-chromatin spaces in the highly organized nucleus and consist of specific factors that function in various nuclear processes (Banani, Lee et al., 2017). ArcRNAs sequestrate sets of specific RNA-binding proteins to initiate building of specific NBs near transcription sites (Chujo et al., 2016, Clemson, Hutchinson et al., 2009). Nuclear stress bodies (nSBs) were primarily reported 30 years ago (Mahl, Lutz et al., 1989) as primate-specific NBs that respond to thermal and chemical stresses. The assembly of nSBs is initiated alongside HSF1-dependent transcription of the primate-specific highly repetitive satellite III (HSATIII) lncRNA (Jolly, Usson et al., 1999) and heat shock-induced HSF1 aggregation (Jolly, Morimoto et al., 1997). HSATIII lncRNAs are transcribed from pericentromeric HSATIII repeated arrays on several human chromosomes (Denegri, Moralli et al., 2002, Jolly, Konecny et al., 2002). These arrays are usually located in heterochromatinic regions that are transcriptionally silent; however, upon exposure to thermal stress, they are rapidly euchromatinized and produce HSATIII lncRNAs (Biamonti & Vourc’h, 2010). HSATIII lncRNAs are retained on chromosomes near their own transcription sites for several hours, even after stress removal, and recruit various RNA-binding proteins, including SAFB, HNRNPM, SRSF1, SRSF7, and SRSF9 (Denegri, Chiodi et al., 2001, Metz, Soret et al., 2004, Weighardt, Cobianchi et al., 1999), as well as specific chromatin-remodeling factors (Hussong, Kaehler et al., 2017, Kawaguchi, Tanigawa et al., 2015) and transcription factors (Jolly, Metz et al., 2004), which results in the assembly of nSBs.

SRSF1, SRSF7, and SRSF9 are serine and arginine-rich (SR) pre-mRNA splicing factors (SRSFs) that contain one or two RNA recognition motifs and a signature RS domain, an intrinsically disordered stretch of multiple SR (in part, SP or SK) dipeptides, near the C-terminus (Long & Caceres, 2009, Shepard & Hertel, 2009). The RS domain is required for protein-protein interactions with other SRSFs or RS domain-harboring proteins (Xiao & Manley, 1997). In addition, the functions of SRSFs in pre-mRNA splicing and mRNA export are modulated through phosphorylation of the RS domain (Cao, Jamison et al., 1997). Notably, the phosphorylation state of the RS domain is dynamically changed in response to environmental transitions, such as thermal stress or circadian changes in body temperature (Guil & Caceres, 2007, Preussner, Goldammer et al., 2017). Upon exposure to thermal stress, protein phosphatase 1 is activated to de-phosphorylate the RS domains of SR proteins, leading to modulation of a wide variety of pre-mRNA splicing events (Shi & Manley, 2007). Concomitantly, CDC2-like kinases 1 and 4 (CLK1 and CLK4), members of the nuclear SR protein kinase family, are induced upon thermal stress exposure to promote re-phosphorylation of SRSFs after stress removal (Ninomiya, Kataoka et al., 2011). These reports raise the intriguing possibility that nSBs are involved in the regulatory network of thermal stress-responsive pre-mRNA splicing by concentrating or sequestering SRSFs; however, the function of nSBs remains poorly investigated.

Here, we performed a genome-wide transcriptomic analysis of HeLa cells following knockdown of HSATIII lncRNAs to deplete nSBs, and found that HSATIII mainly promotes intron retention by hundreds of genes. Further investigation revealed that nSBs accelerate intron retention specifically during recovery from thermal stress. We also explored the protein composition of nSBs using chromatin isolation by Comprehensive identification of RNA-binding proteins by mass spectrometry (ChIRP-MS) and an antisense oligonucleotide (ASO) of HSATIII lncRNA as a stable core component of nSBs. This analysis identified 141 proteins as putative nSB components in HeLa cells. Among them, we focused on CLK1, a SR protein kinase that is specifically recruited to nSBs during thermal recovery to re-phosphorylate SRSFs. Notably, we found that nSBs accelerate the rate of re-phosphorylation of SRSFs after stress removal. Phosphorylated SRSFs then promote retention of the HSATIII target introns. Based on our findings, we propose that nSBs serve as a platform to control the temperature-dependent phosphorylation states of selected SRSFs to regulate intron retention by hundreds of pre-mRNAs. The transcriptomic and subcellular proteomic analyses described here unveil the molecular events occurring in nSBs, which have remained enigmatic for the 30 years since their discovery.

## Results

### HSATIII lncRNAs facilitate intron retention in hundreds of mRNAs during stress recovery

To investigate the role of nSBs in stress-dependent gene expression, particularly splicing regulation, we analyzed global transcriptomic changes in HSATIII-depleted HeLa cells. Defective nSB formation was confirmed in HSATIII ASO-treated (ΔHSATIII) cells exposed to thermal stress at 42°C for 2 h and recovery at 37°C for 1 h (Figure 1A). Subsequently, RNA sequencing (RNA-seq) was used to compare the transcriptomes of control and ΔHSATIII cells. To efficiently detect changes in splicing patterns, nuclear poly(A)+ RNAs were prepared from the isolated nuclei of control and ΔHSATIII cells during the thermal stress recovery phase, and the expression levels of each annotated exon and intron were compared. The expression levels of 533 introns in 434 genes and 3 exons in 3 genes were significantly decreased upon HSATIII knockdown (ΔHSATIII/control<0.5) (Figure 1C and 1D). In addition, the expression levels of 17 introns in 17 genes and 4 exons in 4 genes were significantly increased upon HSATIII knockdown (ΔHSATIII/control>2) (Figure 1C and 1D). For example, HSATIII knockdown reduced the expression levels of introns 2 and 3 of *TAF1D* (Figure 1E, blue bars), and decreased and increased the expression levels of intron 2 (blue bar) and exon 3 (red bar) of *DNAJB9*, respectively (Figure 1F). These findings suggest that HSATIII lncRNAs mainly promote intron retention of pre-mRNAs in thermally stressed cells.

**Figure 1.**
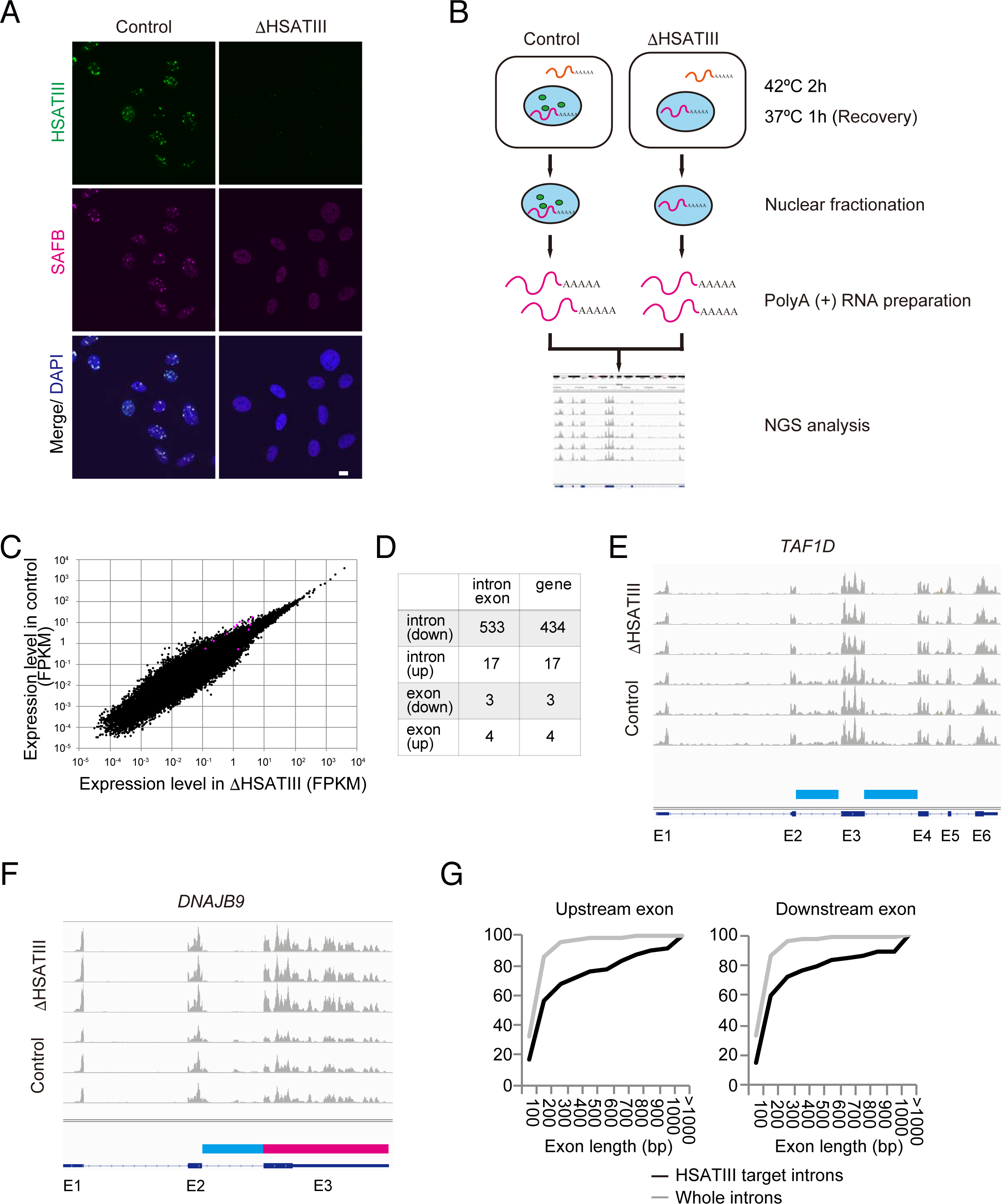
HSATIII lncRNAs control intron retention of a specific set of genes A HSATIII ASO-mediated depletion of nSBs. HeLa cells were transfected with a HSATIII ASO (ΔHSATIII) or HSATIII sense oligonucleotide (control), exposed to thermal stress (42°C for 2 h and recovery for 1 h at 37°C), and then visualized by HSATIII-FISH (green) and immunofluorescence using an anti-SAFB antibody (magenta). The nuclei were stained with DAPI (blue). Scale bar: 10 μm. NGS, next-generation sequencing. B The method used to screen for HSATIII-regulated genes during thermal stress recovery. C The expression levels of all detected introns in control and ΔHSATIII cells (n=3). Magenta dots indicate introns that were experimentally validated in subsequent experiments. D The numbers of introns and exons that were affected by HSATIII knockdown (fold change>2). E, F Examples of RNA-seq read maps of HSATIII target RNAs. Affected introns and exons are indicated by blue (down-regulated upon HSATIII knockdown) and magenta (up-regulated) boxes. G Cumulative frequency curves of the lengths of adjacent exons of 399 HSATIII-up-regulated internal introns. The lengths of whole annotated internal introns are shown as a reference.

We noticed that the sizes of the exons adjacent to HSATIII-targeted introns tended to be longer than the average size of exons (Figure 1G, see also Figure EV4F). To avoid sampling bias caused by the first and the last exons, which tend to be longer than internal exons, we compared the sizes of the exons adjacent to 399 HSATIII-targeted internal introns with those adjacent to 175,923 whole annotated human internal introns. Overall, 8.8% of the exons upstream and 10.3% of the exons downstream of HSATIII-targeted introns were longer than 1 kb (Figure 1G). By contrast, only 0.7% of exons adjacent to the whole annotated internal introns were longer than 1 kb, confirming that the exons adjacent to HSATIII-targeted introns tended to be longer than the average exon size.

A gene ontology analysis of 434 genes in which intron retention was promoted by HSATIII revealed a significant enrichment (FDR<0.05) of genes associated with multiple functions, including DNA/RNA metabolism, biosynthesis, stress response, and cell cycle (Table EV1). Only 57 and 30 of the 533 HSATIII-targeted introns were previously identified as retained introns (also referred to as detained introns) in the human ENCODE database and a human glioblastoma cell line, respectively (Table EV2, discussed later) (Boutz, Bhutkar et al., 2015, Braun, Stanciu et al., 2017).

### HSATIII lncRNAs promote intron retention during thermal stress recovery

To validate the RNA-seq data, we indiscriminately selected 11 introns that were affected by HSATIII knockdown (10 down-regulated and 1 up-regulated; FPKM>0.5 in the control) and used quantitative RT-PCR (qRT-PCR) to analyze their levels in total RNAs from control and ΔHSATIII cells. The levels of the intron-retaining pre-mRNAs were examined under three conditions: normal (37°C), thermal stress at 42°C for 2 h, and thermal stress at 42°C for 2 h followed by recovery at 37°C for 1 h (Figure 2A). For all down-regulated introns examined, the levels of intron retention were comparable in control and ΔHSATIII cells during thermal stress, but were significantly lower in ΔHSATIII cells than in control cells during the recovery phase (Figure 2A, upper panel). Notably, we recognized two distinct classes of intron retention during temperature transition: class 1 introns were retained at substantial levels under normal conditions, excised during thermal stress, and re-accumulated during stress recovery; and class 2 introns were mostly excised under normal and thermal stress conditions, but retained during stress recovery (Figure 2A). Intron 1 of *PFKP* was an exceptional up-regulated intron that was retained during stress recovery of ΔHSATIII cells (Figure 2A). A qRT-PCR analysis of subcellularly fractionated nuclear and cytoplasmic RNAs confirmed that all of the intron-retaining pre-mRNAs mentioned above were retained in the nucleus (Figure 2B), suggesting that mRNA export is prevented by the intron retention. In contrast to the marked effect on the levels of intron-retaining pre-mRNAs, HSATIII knockdown scarcely affected the levels of the cognate intron-removed (spliced) mRNAs. As exceptions, the levels of the spliced *DNAJB9*, *CLK1*, and *PFKP* mRNAs were significantly higher (*CLK1*, *DNAJB9*) or lower (*PFKP*) in ΔHSATIII cells than in control cells (Figure 2A, lower panel).

**Figure 2.**
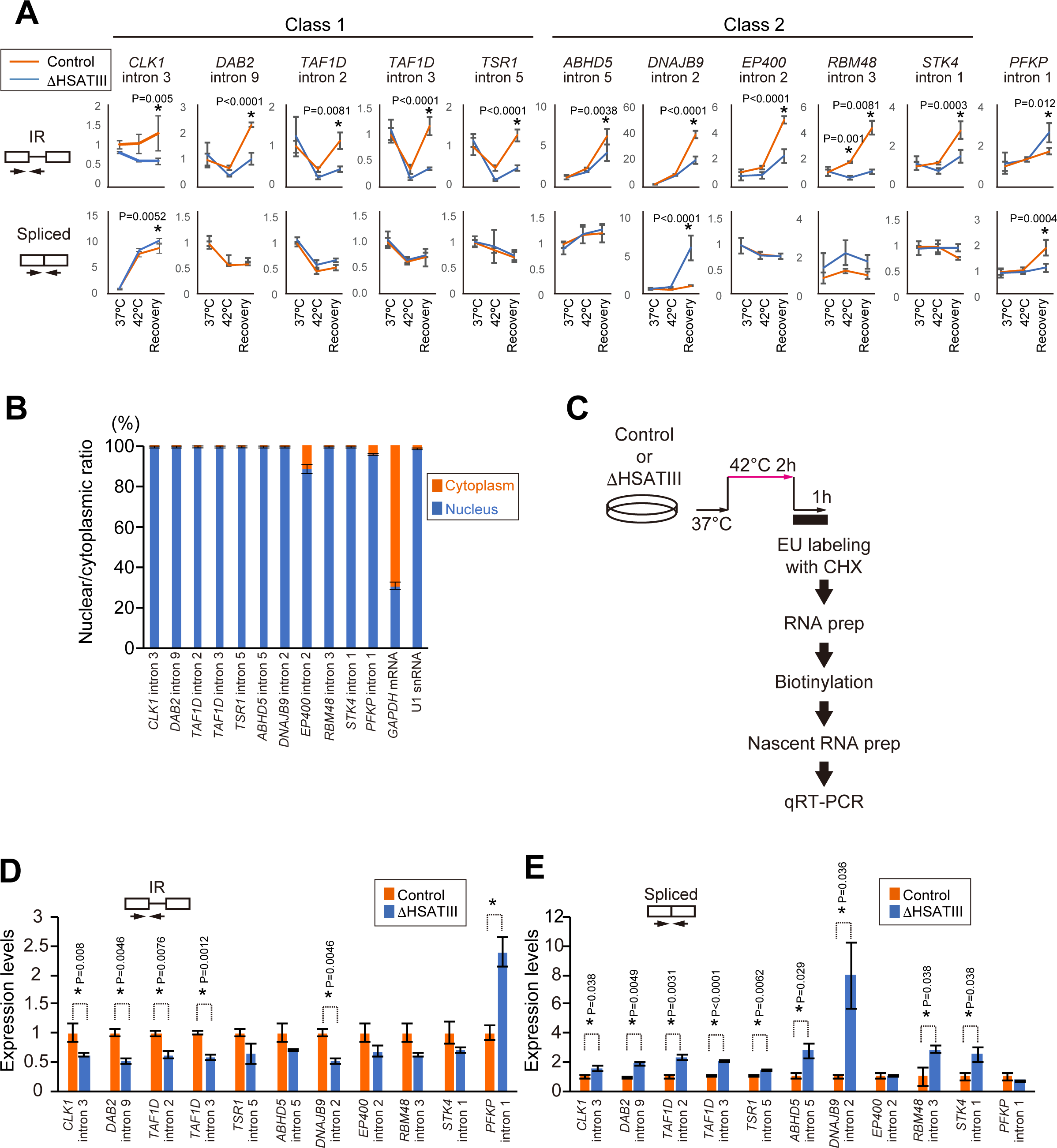
HSATIII lncRNAs control intron retention by regulating splicing. A Validation of HSATIII target introns by qRT-PCR. The graphs show the relative amounts of the intron-retaining (IR) (upper) and spliced (lower) forms in control and HSATIII knockdown cells under three conditions: 37°C (normal), 42°C for 2 h (thermal stress), and thermal stress followed by recovery at 37°C for 1 h. Expression levels were calculated as the ratio of each RNA to *GAPDH* mRNA and were normalized to the levels in control cells under normal conditions (37°C). Data are shown as the mean±SD (n=3); \**p*<0.05 (Sidak’s multiple comparison test). B Nuclear localization of the intron-retaining RNAs. The relative amounts of intron-retaining RNAs in the nuclear and cytoplasmic fractions were quantified by qRT-PCR and are represented as the ratio (% of the total). *GAPDH* mRNA and U1 snRNA were used as cytoplasmic and nuclear controls, respectively. Data are shown as the mean±SD (n=3). C Overview of the qRT-PCR analysis of newly synthesized RNA within 1 h after thermal stress removal. CHX, Cycloheximide; EU, 5-Ethynyl uridine D, E The levels of HSATIII target introns in newly synthesized RNAs within 1 h after thermal stress removal, as determined by qRT-PCR. The graphs show the changes in the expression levels of the intron-retaining (IR) (D) and spliced (E) forms. Expression levels were calculated as the ratio of each RNA to *GAPDH* mRNA and were normalized to the level in the control cells. Data are shown as the mean±SD (n=3); \**p*<0.05 (multiple t-test modified by Holm-Sidak’s method).

Next, the levels of the intron-retaining pre-mRNAs and the spliced mRNAs were measured in newly synthesized nascent RNA pools captured by pulse-labeled RNAs with ethinyl uridine (see Figure 2C and STAR methods). As shown in Figure 2D and 2E, with the exception of *PFKP*, the levels of the intron-retaining pre-mRNAs were lower in ΔHSATIII cells than in control cells. By contrast, with the exceptions of *EP400* and *PFKP*, the levels of the spliced mRNAs were significantly higher in ΔHSATIII cells than in control cells. Overall, these data suggest that HSATIII promotes intron retention by suppressing splicing of newly synthesized transcripts.

### HSATIII affects the kinetics of accumulation of intron-retaining pre-mRNAs during thermal recovery

Among the retained introns regulated by HSATIII, a subpopulation of the *CLK1* mRNA reportedly localizes in the nucleus as a partially unspliced pre-mRNA that retains introns 3 and 4 (Figure 3A) (Duncan, Howell et al., 1995, Ninomiya et al., 2011). Excision of retained introns is induced by thermal and osmotic stresses, neuronal activity, and inhibition of CLK1 kinase activity to produce mature mRNAs (Mauger, Lemoine et al., 2016, Ninomiya et al., 2011). We also detected another spliced isoform of the *CLK1* mRNA produced by skipping of exon 4, which is committed to nonsense-mediated mRNA decay (Figure 3A). Consequently, we examined the effect of HSATIII knockdown on thermal stress-responsive excision of the retained introns of the *CLK1* pre-mRNA at several time points using semi-quantitative RT-PCR. As reported previously (Ninomiya et al., 2011), the retained introns were excised to form the mature mRNA in both control and ΔHSATIII cells after a 2 h thermal stress exposure (Figure 3B, 42°C, 2 h). In control cells, the level of the intron 3 and 4-retaining *CLK1* pre-mRNA was restored within 1 h after stress removal (Figure 3B, lanes 6–10, and Figure 3C). Notably, this process was markedly delayed in ΔHSATIII cells, in which restoration of the original level of the intron 3 and 4-retaining *CLK1* pre-mRNA took longer than 4 h (Figure 3B, lanes 1–5, and Figure 3C).

**Figure 3.**
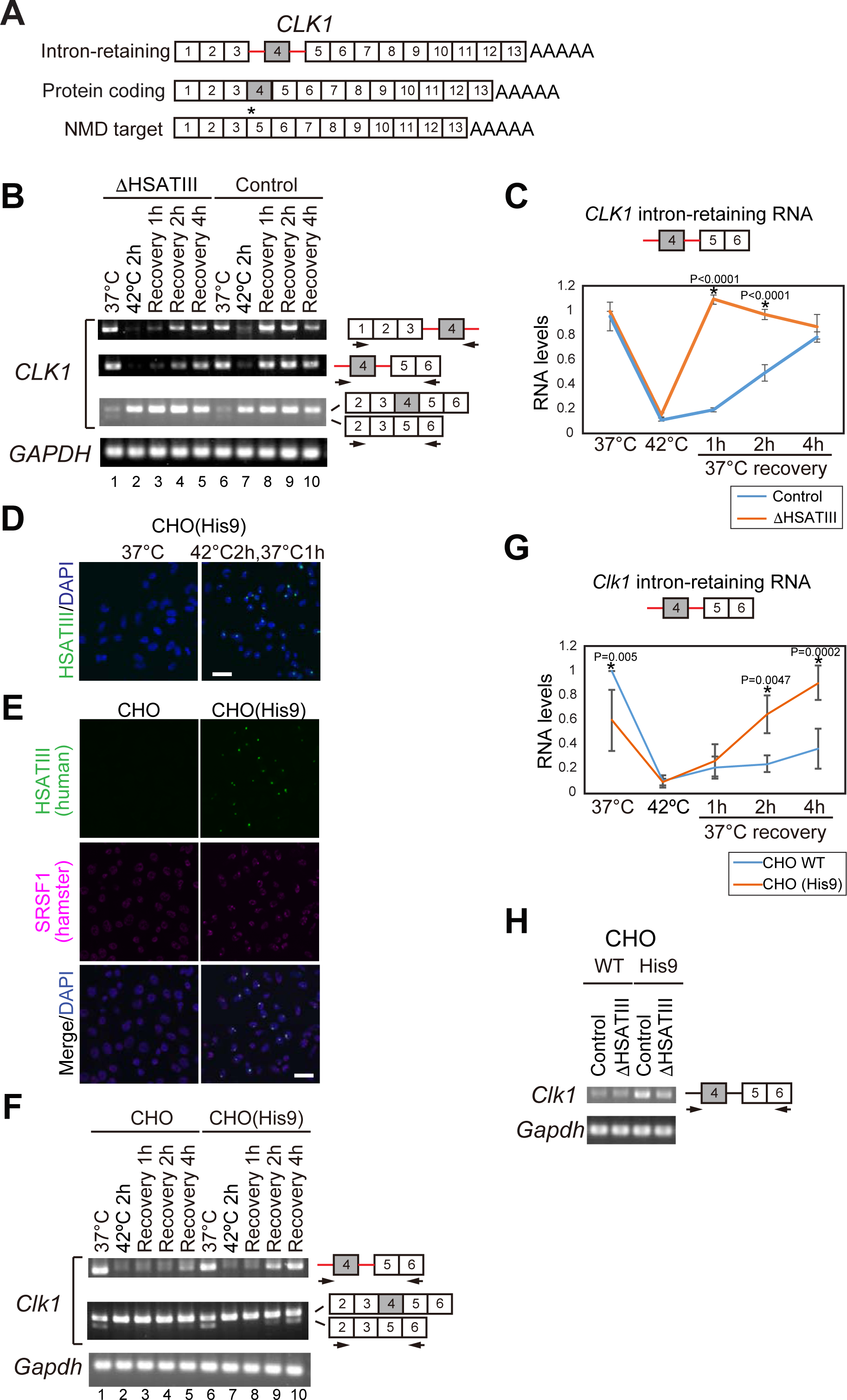
The HSATIII lncRNA is necessary and sufficient to promote nuclear intron retention A Splicing isoforms of the *CLK1* pre-mRNA. The retained introns are indicated by red lines. An asterisk indicates the position of the premature termination codon in the nonsense-mediated mRNA decay-targeted isoform. B Time course analysis of the splicing pattern of *CLK1* pre-mRNAs in control and HSATIII knockdown (ΔHSATIII) cells by semi-quantitative RT-PCR. Arrows indicate the positions of PCR primers. The *GAPDH* mRNA was used as an internal control. C Quantification of the data shown in (B). Data are shown as the mean±SD (n=3); \**p*<0.05 (Sidak’s multiple comparison test). D Thermal stress-induced expression of HSATIII in CHO (His9) cells. The cells were visualized by HSATIII-FISH (green), and the nuclei were stained with DAPI (blue). E Ectopic HSATIII-induced nSB assembly in CHO (His9) cells. The nSBs were visualized by HSATIII-FISH (green) and immunofluorescence using an anti-SRSF1 antibody (magenta). The nuclei were stained with DAPI. Scale bar: 10 μm. F Time course analysis of the splicing pattern of Chinese hamster *Clk1* pre-mRNAs in control (CHO) and CHO (His9) cells by semi-quantitative RT-PCR. Arrows indicate the positions of PCR primers. The *GAPDH* mRNA was used as an internal control. G Quantification of the data shown in (F). Data are shown as the mean±SD (n=3); \**p*<0.05 (Sidak’s multiple comparison test). H The effect of HSATIII knockdown on pooling of intron-retaining RNAs in control (CHO) and CHO (His9) cells (42°C for 2 h and recovery for 1 h at 37°C).

Like *CLK1*, *TAF1D* has two HSATIII target introns (Figure 1E) that border a cassette-type exon 3 (Figure EV1A). Semi-quantitative RT-PCR confirmed that the partially unspliced *TAF1D* pre-mRNA retaining introns 2 and 3 underwent splicing upon thermal stress and was rapidly re-accumulated after stress removal in control cells (Figure EV1B). However, this re-accumulation was markedly delayed in ΔHSATIII cells (Figure EV1B). In addition, we also examined the levels of the *DNAJB9* mRNA, another intron-retaining RNA that harbors a single HSATIII target intron (Figure EV1A), and found that intron retention was detectable in control cells but not ΔHSATIII cells (Figure EV1B). We also confirmed that the thermal stress-responsive alternative splicing of the *HSP105* and *TNRC6a* mRNAs reported previously (Yamamoto, Furukawa et al., 2016, Yasuda, Nakai et al., 1995) was barely affected by HSATIII knockdown (Figure EV1C), indicating selectivity of target pre-mRNAs by HSATIII lncRNAs.

To further support the proposal that HSATIII lncRNAs promote intron retention, we used a gain-of-function approach using a somatic hybrid Chinese hamster ovary (CHO) cell line possessing human chromosome 9 (CHO (His9)) (Tanabe, Nakagawa et al., 2000). Since HSATIII is primate-specific, these lncRNAs are only expressed from human chromosome 9 in CHO (His9) cells upon thermal stress, and form nSB-like granules with hamster SRSF1 (Figure 3D and 3E). A time course analysis of the splicing pattern of the endogenous hamster *Clk1* mRNA revealed that re-accumulation of the intron-retaining pre-mRNA after stress removal was markedly faster in CHO (His9) cells than in control CHO cells (Figure 3F and 3G). Notably, HSATIII knockdown abolished the acceleration effect in CHO (His9) cells (Figure 3H). Taken together, these data suggest that HSATIII lncRNAs are required and sufficient for acceleration of intron retention of the *CLK1* pre-mRNA during stress recovery, and this process was likely acquired in primate species.

### Multiple SRSFs and SR-related proteins interact with HSATIII in nSBs

To examine the molecular mechanism of intron retention by HSATIII lncRNAs, we attempted to comprehensively identify the proteins associated with HSATIII, which likely correspond to nSB components. ChIRP was employed to pull down the ribonucleoprotein (RNP) complexes of HSATIII using a biotinylated ASO (Figure 4A). Because HSATIII lncRNAs consist of multiple GGAAU repetitive sequences (Jarmuz, Glotzbach et al., 2007, Valgardsdottir, Chiodi et al., 2005), HSATIII RNP complexes could be efficiently captured by a single biotinylated HSATIII ASO consisting of four tandem repeats of ATTCC. ChIRP was carried out using formaldehyde-crosslinked HeLa cells that were treated at 42°C for 2 h followed by recovery at 37°C for 1 h. A RT-PCR analysis revealed efficient (>10%) and specific precipitation of HSATIII with the HSATIII ASO from thermally stressed cells (Figure 4B). As a negative control, neither the NEAT1 nuclear lncRNA nor the *GAPDH* mRNA was precipitated with the HSATIII ASO (Figure 4B). Silver staining of the coprecipitated proteins in a SDS-PAGE gel identified multiple bands that were strongly detected in the stressed cells, but were only detected as faint background signals in the control samples (Figure 4C). A liquid chromatography mass spectrometry analysis revealed that 141 proteins, most of which that have not yet been reported as nSB components, were specifically coprecipitated with HSATIII lncRNAs from the stressed cells (Table EV3). Most of these proteins are likely RNA-binding proteins that possess canonical RNA-binding domains. A gene ontology analysis revealed significant enrichment of proteins functionally associated with nuclear RNA processing, such as pre-mRNA splicing, 3’-end processing, and export of mRNAs (Table 1). Notably, most SRSFs and HNRNPs, including known nSB components (SAFB, SRSF1, SRSF7, SRSF9, and HNRNPM), were identified as major components. In addition to SRSFs, non-classical SR proteins and RNA-binding SR-related proteins, which harbor one or more clusters of SR and SP dipeptides, such as TRA2A, TRA2B, U2AF35, U2AF65, LUC7L, SRRM1, BCLAF1, THRAP3, and PPHLN1 (Castello, Fischer et al., 2012, Long & Caceres, 2009), were also identified (Table EV3). Although the number of peptides was limited, the CLK1 SR protein kinase was detected in the ChIRP fraction (described below). By contrast, some previously reported nSB-localized proteins, such as HSF1, CBP, SWI/SNF components, and BRD4, were undetectable in our ChIRP-MS analysis (Table EV3). The nSB localization of the proteins identified above was confirmed by western blotting of the ChIRPed proteins (Figure 4D) and/or immunofluorescence (Figure 4E). The SR-related TRA2B and THRAP3 proteins interacted and colocalized with HSATIII in nSBs (Figure 4D and 4E). The transiently expressed PPHLN1 protein was also colocalized with HSATIII in nSBs (Figure 4E). These results indicate that SR-related proteins, as well as SRSFs, selectively interact with HSATIII in nSBs.

**Figure 4.**
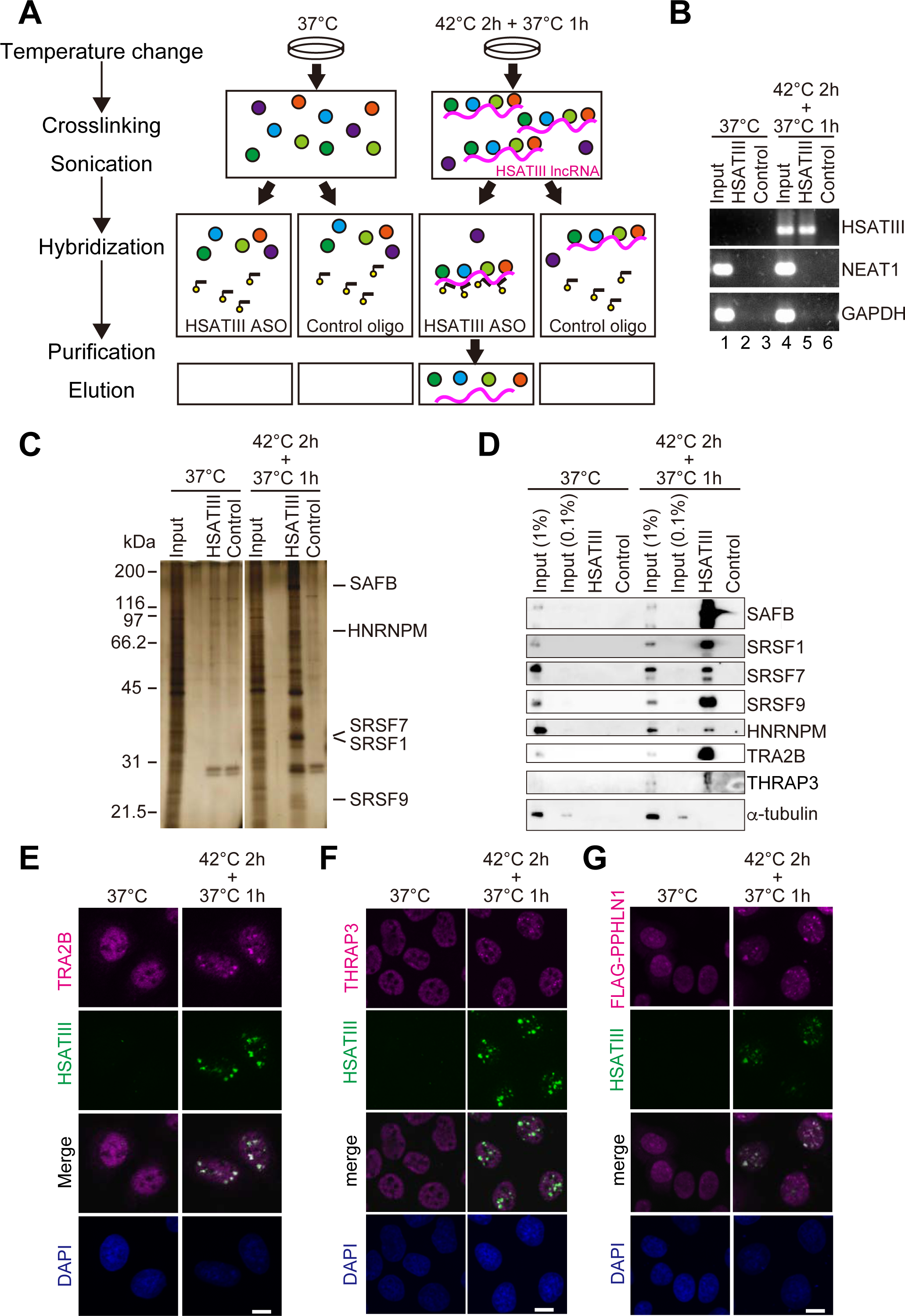
Identification of HSATIII-interacting proteins by ChIRP. A Overview of the ChIRP procedure used to identify HSATIII-interacting proteins in thermal stress-exposed cells. HSATIII-interacting proteins were crosslinked and captured with a biotinylated HSATIII ASO. As a negative control, the same procedure was performed using control (37°C) cells and/or a control probe. B RT-PCR validation of the specific precipitation of HSATIII by ChIRP. The NEAT1 ncRNA and *GAPDH* mRNA were used as negative controls. Input: 100%. C Silver staining of the coprecipitated proteins. The predicted mobilities of five known nSB proteins are indicated. Input: 0.1%. D Western blot analyses of proteins identified by the HSATIII-ChIRP experiment. Known nSB proteins (SAFB, SRSF1, SRSF7, SRSF9, and HNRNPM) were used as positive controls, and α-tubulin was used as a negative control. TRA2B and THRAP3 are newly identified nSB components. E–G Colocalization of novel nSB proteins with HSATIII. Endogenous TRA2B (E), THRAP3 (F), and FLAG-tagged PPHLN1 (G) were stained with anti-TRA2B, anti-THRAP3, and anti-FLAG antibodies, respectively (magenta). HSATIII was visualized by RNA-FISH using a dig-labeled HSATIII ASO (green), and the nuclei were stained with DAPI (blue). Scale bar: 10 μm.

**Table 1.**
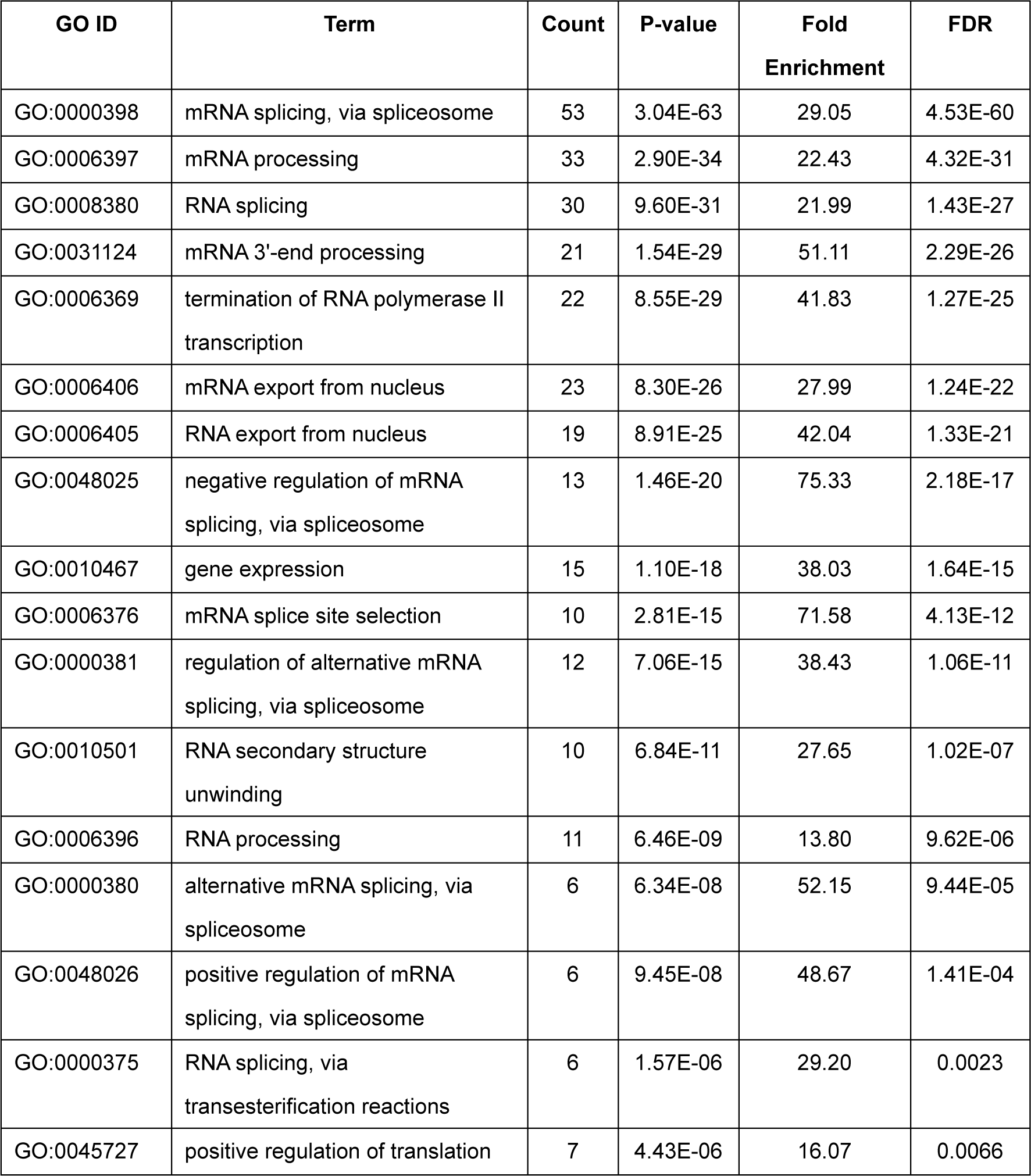
Functional enrichment of HSATIII-interacting proteins

### CLK1 is recruited to nSBs during thermal stress recovery

As described above, CLK1 was detected in the HSATIII-ChIRP fraction (Table EV3). CLK1 is a nuclear SR protein kinase that phosphorylates and modulates the activity of SRSFs in pre-mRNA splicing (Duncan, Stojdl et al., 1997), raising the intriguing possibility that phosphorylation of SRSFs in nSBs is modulated by CLK1. Consequently, we monitored its interaction with HSATIII by ChIRP at five time points during temperature transitions from 37°C to 42°C for 1, 2, or 3 h, or 37°C to 42°C for 2 h followed by recovery at 37°C for 1 h. CLK1 was specifically coprecipitated with HSATIII following recovery after 2 h exposure to thermal stress (Figure 5A). By contrast, CLK1 was poorly detected after thermal stress for 2 or 3 h without recovery at 37°C (Figure 5A), suggesting that the temperature shift from 42°C to 37°C is required to recruit CLK1 onto HSATIII. The nSB components SRSF7 and SRSF9 interacted with HSATIII after a 2 h exposure to thermal stress alone (Figure 5A), suggesting that, unlike CLK1, they do not require the subsequent temperature shift to 37°C for this interaction.

**Figure 5.**
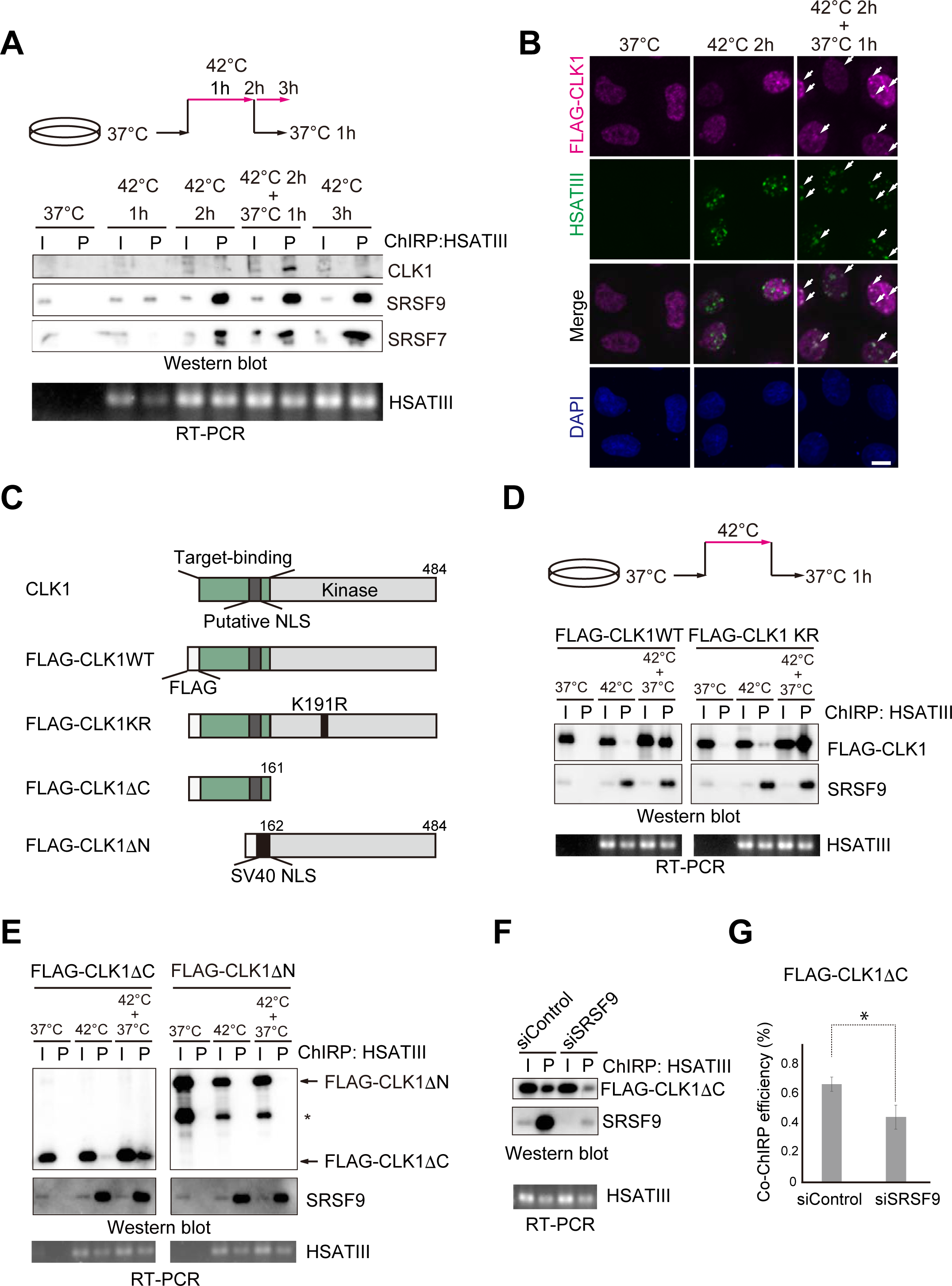
CLK1 is recruited to nSBs during thermal stress recovery A Specific interaction of CLK1 with HSATIII during thermal stress recovery. HSATIII-ChIRP at various time points during and after thermal stress exposure. Endogenous CLK1, SRSF7, and SRSF9 were detected using anti-CLK1, anti-SRSF7, and anti-SRSF9 antibodies, respectively. HSATIII was detected by ChIRP-RNA purification, followed by RT-PCR. Input: 0.1% for CLK1, 1% for SRSF7 and SRSF9, and 100% for HSATIII. B Re-translocation of FLAG-CLK1 to nSBs after thermal stress removal. HeLa cells were transfected with a FLAG-CLK1 expression vector, cultured for 16 h, and then exposed to thermal stress (42°C for 2 h) with or without recovery (37°C for 1 h). FLAG-CLK1 was visualized with an anti-FLAG antibody (magenta). Nuclear stress bodies (green) and nuclei (blue) were visualized by HSATIII-FISH and DAPI staining, respectively. Scale bar: 10 μm. C The domain structures of wild-type (WT) CLK1, the catalytically inactive mutant (CLK1 KR), and the N- and C-terminal partial fragments (CLK1ΔC and CLK1ΔN). D, E Recovery phase-specific interaction of HSATIII with CLK1 proteins. HSATIII-ChIRP/western blotting was performed using HeLa cells expressing FLAG-CLK1 WT or the mutants. FLAG-tagged proteins and SRSF9 were detected by western blotting using an anti-FLAG and anti-SRSF9 antibody, respectively. HSATIII was detected by ChIRP followed by RT-PCR. Input: 1% for western blotting, 100% for RT-PCR. Asterisk: an unidentified partial fragment of FLAG-CLK1ΔN. F Dependency of the CLK1ΔC interaction with HSATIII on SRSF9. HSATIII-ChIRP was performed using control and SRSF9-depleted cells expressing FLAG-CLK1ΔC (42°C for 2 h and recovery for 1 h at 37°C). Input: 1%. G Coprecipitation ratio of CLK1ΔC with HSATIII in the control and SRSF9-depleted cells described in (F). Data are shown as the mean±SD (n=3); \**p*<0.05 (Student’s t-test).

Next, we examined the localization of CLK1 in HeLa cells following thermal stress at 42°C for 2 h and recovery at 37°C for 1 h. We confirmed that transiently expressed FLAG-CLK1 mimicked the behavior of endogenous CLK1 (Figure 5D left). An immunofluorescence analysis revealed that FLAG-CLK1 predominantly localized in nuclear speckles with SRSF2 under normal conditions (Figure EV2A). FLAG-CLK1 was poorly detectable in nSBs after exposure to thermal stress for 2 h, but relocated to nSBs after recovery at 37°C (Figure 5B). These data indicate that the nSB localization of CLK1 occurs during stress recovery and correlates with the interaction with HSATIII lncRNAs.

To obtain mechanistic insights into the temperature-dependent recruitment of CLK1 to nSBs, we used HSATIII-ChIRP to examine the interaction of various CLK1 mutants with nSBs during stress recovery (Figure 5C). The catalytically inactive CLK1 mutant, in which the ATP-binding pocket of the kinase domain was mutated by replacement of lysine 191 with arginine (Flag-CLK1KR) (Colwill, Pawson et al., 1996), interacted with HSATIII and localized to nSBs during stress recovery (Figure 5D, right, and Figure EV2B and EV2C), indicating that the temperature-dependent recruitment of CLK1 to nSBs does not require its catalytic activity. CLK1 interacts with substrate proteins such as SRSFs and homo-dimerizes via its N-terminal intrinsically disordered region (IDR) (Duncan et al., 1995, Keshwani, Hailey et al., 2015). In addition, the C-terminal kinase domain phosphorylates the substrates (Bullock, Das et al., 2009). The C-terminal truncated CLK1 mutant (FLAG-CLK1ΔC in Figure 5C) localized to nuclear speckles under both normal and stressed conditions, and relocated to nSBs after stress removal (Figure EV2D and EV2E); however, the N-terminal truncated CLK1 mutant (FLAG-CLK1ΔN in Figure 5C) was diffusely localized in the nucleoplasm and located to neither nuclear speckles nor nSBs (Figure EV2F and EV2G). The CLK1ΔN mutant possessed a SV40 nuclear localization signal instead of the original NLS present in the deleted N-terminal domain and was properly imported into the nucleus (Figure EV2F–EV2H). CLK1ΔC was coprecipitated with HSATIII much more efficiently under the stress recovery condition than under the thermal stress condition, whereas CLK1ΔN was poorly precipitated under any condition (Figure 5E), indicating that the N-terminal domain of CLK1 is necessary and sufficient for its temperature-dependent interaction with HSATIII. We also examined whether SRSFs anchor CLK1 to nSBs. SiRNA-mediated knockdown of SRSF9, one of the major nSB-localized SRSFs, resulted in a significant reduction in the interaction of CLK1ΔC with HSATIII (Figure 5F and 5G), indicating that SRSF9 contributes to the recruitment of CLK1 at nSBs.

The amino acid sequences of the C-terminal kinase domains of CLK family proteins (CLK1–4) are highly conserved. By contrast, the N-terminal IDRs are similar in CLK1 and CLK4, but are only partly conserved between CLK1/4 and CLK2. Consequently, we examined the involvement of the N-terminal IDRs of ubiquitously expressed CLK2 and CLK4 in nSB recruitment. The N-terminal IDRs of CLK2 and CLK4 were coprecipitated with HSATIII by ChIRP and colocalized with HSATIII after 1 h thermal stress recovery (Figure EV2I and EV2J), indicating that both proteins are able to relocate to nSBs during stress recovery through their poorly conserved N-terminal IDRs.

### nSBs are platforms for the rapid re-phosphorylation of SRSFs by CLK1 during thermal recovery

Next, we examined the subnuclear localization of CLK1 and SRSFs during thermal stress and recovery. First, we confirmed that SRSF1 and SRSF9 colocalized with HSATIII in nSBs after 2 h thermal stress exposure and after stress removal (Figure 6A and Figure EV3A), which is consistent with the ChIRP results (Figure 5A). We also confirmed that the SRSF9 foci were diminished and diffused throughout the nucleoplasm in ΔHSATIII cells (Figure EV3B), and did not overlap with SRSF-rich nuclear speckles (Figure 6B). Moreover, we confirmed colocalization of CLK1 and SRSF9 in nSBs during stress recovery (Figure 6C). SRSFs are rapidly de-phosphorylated upon thermal stress (Shi & Manley, 2007); therefore, it is possible that CLK1 is recruited to nSBs during stress recovery to promote re-phosphorylation of SRSFs. To analyze the phosphorylation states of SRSFs after thermal stress and recovery, we examined SRSF9, which possesses a short RS domain with four SRs and three SPs. Western blotting followed by Phos-tag SDS-PAGE (Kinoshita, Kinoshita-Kikuta et al., 2008) revealed multiple bands corresponding to different phosphorylation states of SRSF9 (Figure 6F). Under normal conditions (37°C), five discrete SRSF9 bands were detected (Figure 6F, lane 1). After thermal stress at 42°C for 2 h (Figure 6F, lane 3), two fast migrating bands (likely hypo-phosphorylated SRSF9) appeared and the upper bands (likely hyper-phosphorylated SRSF9) were diminished (Figure 6F, indicated by arrowheads and arrows, respectively). After recovery at 37°C for 1 h, the band pattern was mostly restored to its original state (Figure 6F, lane 4); however, this restoration was abolished by administration of the CLK1 inhibitor KH-CB19 (Fedorov, Huber et al., 2011) at the beginning of the stress recovery phase (Figure 6D and 6F, lane 5). These findings suggest that the observed band shift to lower positions after heat stress and subsequent restoration after stress removal were caused by thermal stress-induced de-phosphorylation and CLK1-dependent re-phosphorylation of SRSF9, respectively. To determine whether nSBs affect CLK1-dependent re-phosphorylation of SRSF9, we examined the Phos-tag band pattern of SRSF9 in ΔHSATIII and control cells (Figure 6E). Restoration of the band pattern of SRSF9 after stress removal was markedly delayed in ΔHSATIII cells (Figure 6F, right). The hyper-phosphorylated band (Figure 6F, arrow a) was mostly restored after 1 h recovery in control cells, but this restoration was delayed until 4 h after stress removal in ΔHSATIII cells. In addition, the hypo-phosphorylated bands (Figure 6F, arrowheads c and d) were rapidly diminished after 1 h recovery in control cells but this process was delayed in ΔHSATIII cells.

**Figure 6.**
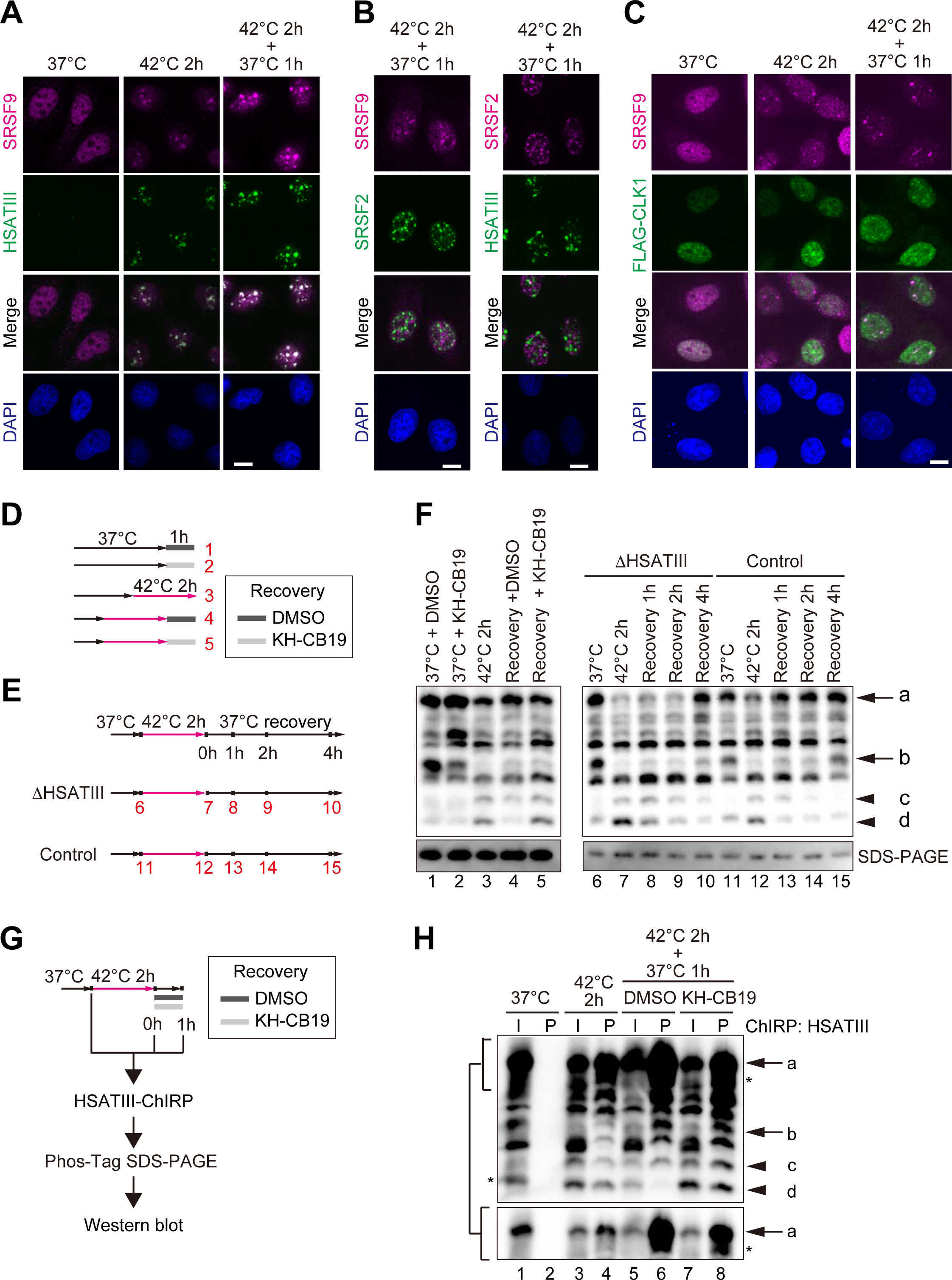
The HSATIII lncRNA accelerates CLK1-dependent re-phosphorylation of SRSF9 A Localization of SRSF9 with HSATIII during thermal stress and recovery. B The subnuclear distribution of nSBs and nuclear speckles. HeLa cells (42°C for 2 h and 37°C for 1 h) were stained with an anti-SRSF9 antibody, an anti-SRSF2 antibody, and a HSATIII-FISH probe. The nuclei were stained with DAPI (blue). C Recovery phase-specific colocalization of CLK1 and SRSF9. HeLa cells were transfected with a FLAG-CLK1 expression vector, cultured for 16 h, and exposed to thermal stress (42°C for 2 h) followed by recovery (37°C for 1 h). FLAG-CLK1 (green) and SRSF9 (magenta) were co-stained using anti-FLAG and anti-SRSF9 antibodies, respectively, and the nuclei were stained with DAPI (blue). Scale bar: 10 μm. D Scheme of sampling references for analysis of the phosphorylation states of SRSF9 in (F). The CLK1 inhibitor KH-CB19 (10 μM) or DMSO as a control was administrated during the recovery period. The numbers correspond to the lane numbers in (F). E Conditions tested in the time course analysis of the phosphorylation states of SR proteins. HeLa cells were transfected with a HSATIII ASO or control SO, cultured for 16 h, exposed to thermal stress (42°C for 2 h), and cultured at 37°C for the indicated period. F The effect of HSATIII knockdown on time course-dependent changes in SRSF9 phosphorylation. The phosphorylation states were estimated by Phos-tag SDS-PAGE, followed by western blotting using an anti-SRSF9 antibody. The arrows and arrowheads indicate the normal phosphorylated forms and thermal stress-induced hypo- and de-phosphorylated forms, respectively. G Scheme of Phos-tag western blotting coupled with HSATIII-ChIRP used to validate the phosphorylation states of SRSF9 within nSBs. H The re-phosphorylation status of SRSF9 within nSBs. Phosphorylated (arrows) and hypo- and de-phosphorylated (arrowheads) SRSF9 in the nSB fractions (lanes 4, 6, and 8) were analyzed by western blotting on Phos-tag gels. KH-CB19 (10 μM) or DMSO as a control was administrated during the recovery period. The lower panel shows a shorter exposure image of the phosphorylated forms. The asterisks indicate bands different from band (a) or (d). Input: 5%.

To confirm that CLK1-dependent re-phosphorylation of SRSF9 occurs within nSBs, we performed Phos-tag western blotting followed by HSATIII-ChIRP analyses of HeLa cells exposed to control conditions (37°C), 42°C for 2 h, or 42°C for 2 h followed by recovery for 1 h at 37°C (Figure 6G and 6H). In the HSATIII-ChIRP fractions, de-phosphorylated SRSF9 (the lower band marked by arrowhead d) was diminished and phosphorylated SRSF9 (marked by arrow a) was markedly increased during the 1 h recovery (Figure 6H, lanes 2, 4, 6, and 8). Notably, these changes were partially inhibited in the presence of KH-CB19 (Figure 6H, lane 8). Western blot analyses of the HSATIII-ChIRP samples revealed that the amount of coprecipitated SRSF9 did not change markedly during the 1 h recovery, regardless of the presence or absence of KH-CB19 (Figure EV3C). Taken together, these data suggest that CLK1 recruited during stress recovery re-phosphorylates SRSF9 within nSBs. Although differentially phosphorylated bands were not clearly separated on the Phos-tag gel, stress-induced mobility shifts of nSB-localized SRSF1 and SRSF7 were restored after stress removal in a CLK1-dependent manner (Figure EV3D), suggesting that CLK1 rapidly re-phosphorylates various SRSFs, and probably SR-related proteins, in nSBs during stress recovery.

### CLK1-dependent phosphorylation of SRSF9 in nSBs underlies the promotion of intron retention during stress recovery

The findings described above raised the intriguing possibility that recruitment of CLK1 to nSBs for rapid re-phosphorylation of SRSFs promotes intron retention of mRNAs during thermal stress recovery. To address this possibility, we investigated the effect of SRSF knockdown on intron retention. Depletion of SRSF7 or SRSF9 barely affected nSB integrity; however, depletion of SRSF1 remarkably disrupted nSBs (Figure EV4A and EV4B), such that we were unable to investigate the effect of SRSF1 on nSB-dependent splicing regulation. Consequently, we investigated the effect of SRSF9 knockdown on intron retention of HSATIII target mRNAs using newly synthesized nascent RNA pools (Figure 7A). With the exception of *PFKP*, SRSF9 knockdown reduced the levels of the intron-retaining mRNAs (Figure 7A) and increased the levels of the spliced forms of the mRNAs (Figure 7B). These results suggest that SRSF9 suppresses pre-mRNA splicing to promote HSATIII-dependent intron retention. RT-PCR analyses revealed that SRSF9 depletion diminished the prominent accumulation of the *CLK1* and *TAF1D* mRNAs retaining two sequential introns after 1 h recovery from thermal stress (Figure 7C and 7D). Consistently, publicly available eCLIP data showed that multiple eCLIP tags of SRSF9 mapped to the exons adjacent to the retained introns of the *CLK1* and *TAF1D* mRNAs in HepG2 cells (Figure EV4C).

**Figure 7.**
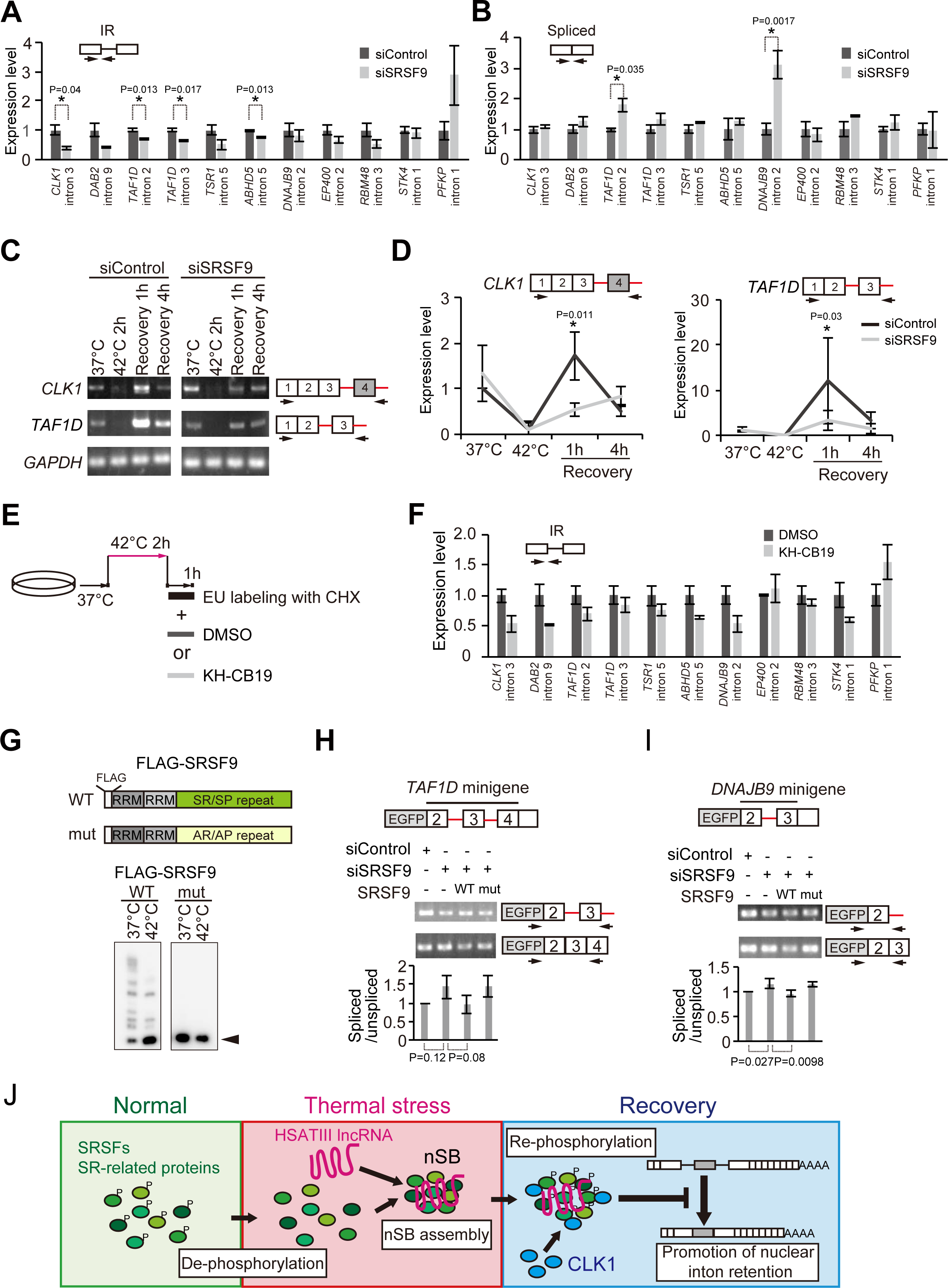
The role of SRSF9 re-phosphorylation in controlling HSATIII-dependent intron retention A, B Validation of the effect of SRSF9 on retention/splicing of HSATIII target introns by qRT-PCR. The relative expression levels of intron-retaining (IR) forms (A) and spliced forms (B) of newly transcribed RNAs during the recovery period are shown. See also the scheme in Figure 2B. Expression levels were calculated as the ratio of each RNA to *GAPDH* mRNA and were normalized to the control. Data are shown as the mean±SD (n=3); \**p*<0.05 (multiple t-test modified by Holm-Sidak’s method). C Time course analysis of the splicing patterns of *CLK1* and *TAF1D* intron-retaining RNAs in control and HSATIII knockdown cells by semi-quantitative RT-PCR. Arrows indicate the positions of primers for RT-PCR. *GAPDH* mRNA was used as an internal control. The experiment was performed in triplicate. D Quantification of the data shown in (C); \**p*<0.05 (Sidak’s multiple comparison test). E Scheme for analyzing the effect of CLK1 inhibition on intron retention by newly transcribed RNAs after stress removal. CHX, Cycloheximide; EU, 5-Ethynyl uridine. E Validation of the requirement of CLK1 catalytic activity for promotion of intron retention by qRT-PCR. F The diagrams show the domain structures of wild-type (WT) and mutant (mut) SRSF9, in which all SR/SP dipeptides were replaced by AR/AP. FLAG-SRSF9-WT and FLAG-SRSF9-mut were extracted from transfected HeLa cells under normal (37°C) and thermal stress (42°C for 2 h) conditions, separated in a Phos-tag gel, and detected by western blot using an anti-FLAG antibody. The arrowhead indicates the thermal stress-induced de-phosphorylated form of FLAG-SRSF9-WT. H, I The role of phosphorylation of SRSF9 on splicing of the *TAF1D* (H) *and DNAJB9* (I) reporters. Each *s*plicing reporter plasmid was transfected into HeLa cells along with a siRNA (control or SRSF9-specific) and a siRNA-resistant SRSF9-WT or SRSF9-mut expression plasmid (or pEGFP-C1 vector as the empty vehicle (-)) 48 h before the assay. Semi-quantitative RT-PCR analyses revealed the splicing patterns of the reporter RNAs purified from the thermal stress-exposed cells (42°C for 2 h and 37°C for 1 h). The positions of the PCR primers are indicated by arrows. Graphs represent the relative changes in the spliced/unspliced ratios, normalized to the control sample with control siRNA and empty plasmid (-). Data are shown as the mean±SD (n=3); \**p*<0.05 (Dunnett’s multiple comparison test). J A model for the function of nSBs in splicing control. Thermal stress-induced HSATIII integrates de-phosphorylated SRSFs and SR-related proteins. After stress removal, CLK1 relocates to nSBs to re-phosphorylate SRSFs, especially SRSF9. Re-phosphorylated SRSF9 suppresses splicing of target introns to accumulate intron-retaining RNAs in the nucleus.

Next, we examined the possibility that CLK1-dependent re-phosphorylation of SRSF9 in nSBs is responsible for the regulation of intron retention. To investigate its requirement for the changes in intron retention after stress removal, CLK1 kinase activity was impeded by KH-CB19 during thermal stress recovery (Figure 7E and 7F). HSATIII-dependent intron retention (and the opposite effect on the *PFKP* mRNA) was impaired in the presence of KH-CB19 (*EP400* was an exception), which supports our hypothesis that CLK1-dependent re-phosphorylation of SRSFs is required for the promotion of intron retention. To directly investigate the role of phosphorylation of SRSF9 in splicing regulation, we compared the effects of wild-type SRSF9 (SRSF9-WT) and its de-phosphorylated mutant (SRSF9-mut), in which all SP and SR dipeptides were replaced by AP and AR, respectively (Figure 7G and Figure EV4D), on splicing of the *TAF1D* and *DNAJB9* splicing reporter minigenes. Western blot analyses of a Phos-tag gel detected multiple FLAG-SRSF9-WT protein bands that shifted to lower hypo- or de-phosphorylated bands after thermal stress (Figure 7G). By contrast, the SRSF9-mut was detected as a single band with the same mobility as the thermal stress-induced de-phosphorylated WT protein, even at 37°C (Figure 7G, arrowhead), and was no longer shifted by thermal stress (Figure 7G), indicating that the SRSF9-mut mimics de-phosphorylated SRSF9.

To replace endogenous SRSF9 with FLAG-SRSF9-WT or the de-phosphorylated mutant, HeLa cells were co-transfected with SRSF9 siRNA and siRNA-resistant FLAG-SRSF9 expression plasmids (Figure EV4E). These cells were also transfected with splicing reporter *TAF1D* and *DNAJB9* plasmids, and the splicing patterns were examined after exposure to thermal stress for 2 h followed by recovery for 1 h (Figure 7H and 7I). SRSF9-WT suppressed splicing of the reporter RNAs, whereas SRSF9-mut did not, suggesting that phosphorylation of the RS domain of SRSF9 is required to promote intron retention of the *TAF1D* and *DNAJB9* mRNAs by suppressing splicing. Taken together, these findings suggest that the re-phosphorylation of the RS domain of SRSF9 by CLK1 in nSBs promotes intron retention of target mRNAs during thermal stress recovery.

## Discussion

In this study, next-generation sequencing-based screening of the regulatory targets of nSBs identified a major molecular function of nSBs in promoting intron retention by more than 400 mRNAs during thermal stress recovery. In recent years, a number of studies have examined conditional intron retention in the nucleus and its biological functions, such as regulation of stress responses, the cell cycle, neuronal activity, learning and memory, spermiogenesis, and tumor growth (Boutz et al., 2015, Braun et al., 2017, Dominguez, Tsai et al., 2016, Gill, Park et al., 2017, Mauger et al., 2016, Naro, Jolly et al., 2017). Thus, nuclear intron retention is being accepted as a regulatory mechanism that fine tunes gene expression under various biological conditions. Here, we found that nSBs constructed with HSATIII arcRNA scaffolds are novel molecular machineries that regulate nuclear intron retention during thermal stress recovery. Comparison of the 533 retained HSATIII-targeted introns identified here with the 3968 retained introns previously reported in the U87 human glioblastoma cell line revealed an overlap of only 30 introns (Table EV2). It is possible that different introns are selected for retention under different physiological conditions, presumably through the use of distinct regulatory factors.

We confirmed that SRSF9 promotes intron retention of HSATIII-targeted pre-mRNAs (Figure 7A). SRSFs are generally considered to promote splicing in a phosphorylation-dependent manner; however, SRSF9 is a repressor of 3’ splice site utilization (Simard & Chabot, 2002). In this case, SRSF9 suppresses splicing by recognizing a specific intronic CE9 element; however, no sequence motif similar to CE9 was found in the HSATIII-dependent retained introns that we identified. According to the public eCLIP database, multiple peaks of SRSF9 eCLIP tags were detectable in the HSATIII-targeted retained introns, as well as adjacent upstream and downstream exons; however, no significant enrichment of SRSF9 eCLIP tags compared with the average numbers in general introns and exons was detected (Figure EV4F), suggesting that SRSF9 is required but not sufficient for HSATIII-dependent intron retention.

Consistent with previous reports (Boutz et al., 2015, Ninomiya et al., 2011), we found that the catalytic activity of CLK1 is required for HSATIII-dependent intron retention. We also found that CLK1 phosphorylates SRSF9 during thermal stress recovery; this phosphorylation was detected in the HSATIII-ChIRP fraction, suggesting that it occurs within nSBs. Furthermore, the results of a splicing reporter assay using model constructs for intron retention support the proposal that CLK1-dependent phosphorylation of the serine residues of SR/SP repeats in SRSF9 promotes intron retention (Figure 7G–I). Among the tested introns, the retention of *PFKP* intron 1 was exceptionally down-regulated by HSATIII, SRSF9, and the catalytic activity of CLK1, suggesting that a common molecular mechanism can have opposing effects on a minor population of target introns. Intron retention of *EP400* moderately required SRSF9 but not CLK1 activity (Figure 7A and 7F), suggesting the existence of a CLK1-independent mechanism to promote intron retention. We propose a model of nSB function (Figure 7J) in which SRSFs are globally de-phosphorylated upon thermal stress at 42°C (Shin, Feng et al., 2004), and HSATIII arcRNAs are simultaneously synthesized to form nSBs that sequestrate the de-phosphorylated SRSFs. Under this condition, the retention of class 1 introns is canceled to produce mature mRNAs. Following recovery to normal conditions (37°C), CLK1 is recruited to nSBs, where it re-phosphorylates the sequestrated SRSFs, which then act to rapidly promote retention of class 1 and 2 introns within 1 h (Figure 7J). These intron retention events may occur outside of nSBs since the intron-retaining pre-mRNAs were barely detected in the HSATIII-ChIRP fraction. Therefore, the re-phosphorylated SRSFs likely relocate from nSBs to the nucleoplasm, which is analogous to the mechanism of mobilization of SRSFs from nuclear speckles to the nucleoplasm through phosphorylation by CLK1 (Colwill et al., 1996). Consequently, we argue that nSBs act as a platform for phosphorylation of sequestrated SRSFs by CLK1 to promote intron retention during thermal stress recovery.

Recruitment of CLK1 to nSBs in response to a temperature change from 42°C to 37°C is considered a critical step for rapid re-phosphorylation of SRSFs within nSBs. This temperature-sensitive recruitment of CLK1 requires its N-terminal IDR. CLK family kinases commonly contain long flexible N-terminal IDRs that resemble the RS domains of SRSFs. The N-terminal IDR of CLK1 contacts both the kinase domain and the RS domain of SRSF1 to induce hyper-phosphorylation of SRSF1 (Aubol, Plocinik et al., 2014). We observed that recruitment of CLK1 to nSBs requires the presence of SRSF9, suggesting that the analogous interaction between the CLK1 N-terminal IDR and SRSF9 is responsible for recruitment of CLK1 to nSBs. The mechanism by which the CLK1 N-terminal IDR senses the temperature change remains elusive. It is possible that thermal stress induces a structural change in the N-terminal IDR of CLK1 that interferes with its contact with the RS domains of SRSF9. The N-terminal IDRs of CLK2 and CLK4 also associate with nSBs in a similar temperature-sensitive manner, suggesting that other CLK family kinases contribute to the phosphorylation of SRSFs in nSBs. The amino acid sequences of the N-terminal IDRs are poorly conserved among CLK family kinases, suggesting that short stretches and/or the compositional property of amino acids among the IDRs might be responsible for the temperature-sensitive recruitment of CLKs to nSBs. CLKs are localized in nuclear speckles at normal temperatures, where they phosphorylate SRSFs and SR-related proteins (Colwill et al., 1996, Gui, Lane et al., 1994). The localization of CLK1 to nuclear speckles is maintained under thermal stress, suggesting that the mechanism used to localize CLK1 to nSBs differs from that used for nuclear speckle localization. This variation may be due to the different compositions of SRSFs in nSBs and nuclear speckles.

In addition to classical SRSFs, we identified other SR-related proteins, including TRA2B and BCLAF1/THRAP3, that also have SR/SP dipeptides as potential sites of phosphorylation by CLKs (Aubol, Plocinik et al., 2013). These SR-related proteins reportedly form a complex with SRSF1, SRSF9, SAFB, and CLKs (Nayler, Stratling et al., 1998, Tai, Geisterfer et al., 2003), raising the possibility that multiple SR and SR-related proteins interact through the IDRs to form the massive membrane-less structure of nSBs, where they are collectively re-phosphorylated by CLKs. The effect of HSATIII on the intron retention of *CLK1* and *TAF1D* pre-mRNAs was limited to within 4 h after stress removal (Figure 3B, 3C, and Figure EV1B). This time course of intron retention correlates with that of SRSF9 re-phosphorylation (Figure 6F). It is likely that nSBs function to rapidly fix the proper levels of intron-retaining pre-mRNA in the nucleus during stress recovery. In addition to the role of intron retention as a regulatory mechanism for fine tuning gene expression, it was recently reported that a subset of introns act as ncRNAs that trap the spliceosome and decrease global splicing upon nutrient depletion in yeast (Morgan, Fink et al., 2019, Parenteau, Maignon et al., 2019). In addition, a role of introns in prevention of R-loop and DNA damage accumulation has also been reported (Bonnet, Grosso et al., 2017). Further investigations will reveal the biological significance of HSATIII-dependent intron retention to cellular and physiological events. Multiple proteins associated with other RNA processing events, including 3’-end processing, nuclear export, stability control, and modification of mRNAs (Table 1 and Table EV3), are found in nSBs, suggesting the involvement of nSBs in controlling mRNA processing events other than splicing during thermal stress and recovery. Further elucidation of the cellular and physiological roles of HSATIII-dependent nSBs will give important clues to understand the significance of the satellite repeat expansion that specifically occurs in primate genomes.

## Materials and Methods

### Cell culture

HeLa cells were cultured in Dulbecco’s modified Eagle’s medium (DMEM) (Nacalai Tesque) supplemented with antibiotics (100 U/ml streptomycin and 100 μg/ml penicillin; Sigma) and 10% fetal calf serum (Sigma) at 37°C in 5% CO_2_. CHO cells were cultured in DMEM/Ham’s F-12 medium (Nacalai Tesque). CHO (His9) cells (JCRB Cell Bank) were grown in medium containing 10 mM L-histidinol (Sigma) to maintain human chromosome 9, and cultured in normal medium from the day before the analysis. For thermal stress induction, the cells were incubated at 42°C in an incubator with 5% CO_2_.

### Plasmid construction

To construct plasmids expressing FLAG-tagged full-length and partial proteins, each DNA fragment was amplified from human cDNA by PCR and introduced in-frame into the pcDNA5/FRT/TO-FLAG vector. The FLAG-SRSF9 expression plasmid (pcDNA3-FLAG-SRSF9) was kindly provided by Dr. Kataoka (Univ. Tokyo). Inverted PCR-based mutagenesis was used to construct plasmids expressing the CLK1 KR mutant or the SRSF9 mutant and its siRNA-resistant mutants. To construct the splicing reporter plasmids, *TAF1D* and *DNAJB9* intron-containing DNA fragments were amplified by PCR from human genomic DNA and introduced in-frame into the pEGFP-C1 vector.

### ChIRP

ChIRP was performed as described previously (Chu, Quinn et al., 2012). Cultured cells at 80–90% confluence (one or two 10 cm culture dishes for each sample) were immediately crosslinked in 10 ml of 1% paraformaldehyde (PFA)/PBS per 10 cm dish for 15 min at room temperature, and the crosslinking reaction was stopped by adding 1 ml of 1.5 M glycine. The crosslinked cells were then washed with 10 ml of ice-cold PBS twice, suspended in 1 ml of ice-cold PBS using a scraper, and collected in microtubes by microcentrifugation. After removing the PBS, the cells were resuspended in 350 μl of lysis buffer (50 mM Tris-Cl pH 7.0, 10 mM EDTA, 1% SDS, 1 mM PMSF, cOmplete™ Protease Inhibitor Cocktail (Sigma), and RNAse inhibitor (Thermo Fisher Scientific)) and disrupted using a Bioruptor (Cosmo Bio) (15 sec ON, 45 sec OFF, 118 cycles). Subsequently, the insoluble fraction was removed by microcentrifugation for 10 min at 15°C, and the supernatant was used for ChIRP. Two volumes of hybridization buffer (750 mM NaCl, 1% SDS, 50 mM Tris-Cl pH 7.0, 1 mM EDTA, 15% (v/v) formamide, 1 mM PMSF, cOmplete™ Protease Inhibitor Cocktail (Sigma), and RNAse inhibitor) and the HSATIII antisense or control oligonucleotide were added. The mixture was incubated at 37°C for 4 h, mixed with Dynabeads MyOne Streptavidin C1 (Thermo Fisher Scientific), and incubated for 30 min. The beads were then washed five times for 5 min at 37°C with 0.8 ml of wash buffer (2× SSC and 0.5% SDS). The bound proteins were eluted and reverse-crosslinked by boiling in SDS sample buffer (95°C for 30 min). For mass spectrometry, the bound proteins were eluted with 100 mM Tris-HCl (pH 8.0) and 2% SDS. For silver staining, the eluted proteins were separated by SDS-PAGE and stained using the SilverQuest Silver Staining Kit (Thermo Fisher Scientific) according to the manufacturer’s manual. For Phos-tag western blotting of ChIRP samples, the eluate and input proteins were collected by trichloroacetic acid precipitation and resuspended in new SDS sample buffer. The bound RNAs were eluted by protease K treatment (50°C for 45 min) followed by heating (95°C for 10 min) and purification as described below.

### Mass spectrometry

To remove the SDS from eluted samples, the methanol-chloroform protein precipitation method was used. Briefly, four volumes of methanol, one volume of chloroform, and three volumes of water were added to the eluted sample and mixed thoroughly. The samples were centrifuged at 20,000 g for 10 min, the water phase was carefully removed, four volumes of methanol were added, and the samples were centrifuged at 15,000 rpm for 10 min. Subsequently, the supernatant was removed and the pellet was washed once with 100% ice-cold acetone. The precipitated proteins were re-dissolved in guanidine hydrochloride, reduced with Tris(2-carboxyethyl)phosphine hydrochloride, alkylated with iodoacetamide, and then digested with lysyl endopeptidase and trypsin. The peptide mixture was applied to a Mightysil-PR-18 (Kanto Chemical) frit-less column (45 × 0.150 mm ID) and separated using a 0–40% gradient of acetonitrile containing 0.1% formic acid for 80 min at a flow rate of 100 nL/min. The eluted peptides were sprayed directly into a mass spectrometer (Triple TOF 5600+; AB Sciex). MS and MS/MS spectra were obtained using the information-dependent mode. Up to 25 precursor ions above an intensity threshold of 50 counts/s were selected for MS/MS analyses from each survey scan. All MS/MS spectra were searched against protein sequences of the RefSeq (NCBI) human protein database (RDB) using the Protein Pilot software package (AB Sciex), and decoy sequences were then selected with a false discovery rate <1%.

### Nucleus/cytoplasm fractionation

Nucleus/cytoplasm fractionation was performed as described previously (Mili, Shu et al., 2001), with minor modifications. Cultured cells on a 35 mm dish were suspended in 200 μl of RSB-100 (10 mM Tris-HCl pH 7.4, 100 mM NaCl, and 2.5 mM MgCl_2_) containing 40 μg/ml digitonin, and centrifuged at 2,000 g for 2 min at 4°C. The supernatant was collected as the cytoplasmic fraction. The pellet (nuclear fraction) was resuspended in 200 μl of RSB-100 buffer. The fractions were lysed in 600 μl of TRI Reagent LS for RNA analysis or 200 μl of 2× SDS sample buffer (200 mM dithiothreitol, 100 mM Tris-HCl pH 6.8, 4% SDS, and 20% glycerol) for western blot analysis.

### Transfection and knockdown

For transfection followed by cell staining or ChIRP, plasmids were transfected into cells using TransIT-LT1 reagent (Mirus) 16–18 h before the assay (0.2 μg of plasmid, 100 μl of opti-MEM, and 1 μl of TransIT reagent were used per 1 ml of culture medium). For siRNA knockdown, Stealth siRNA (Thermo Fisher Scientific) was transfected at a final concentration of 10 nM using Lipofectamine RNAiMAX (Thermo Fisher Scientific) 48 h before the experiments. For the splicing reporter assay, the cells were transfected using 12 pmol of Stealth siRNA, 50 ng of splicing reporter plasmid, 300 ng of SRSF9-expressing plasmid, 200 μl of opti-MEM, and 2 μl of Lipofectamine 2000 reagent (Thermo Fisher Scientific) per 1 ml of culture medium, and cultured for 48 h. For nuclear RNA knockdown, a modified antisense or control oligonucleotide (1 μM) was transfected into 3 × 10^6^ cells using Nucleofector technology (Lonza) 16–18 h before the assay. ASOs and siRNAs are listed in Appendix Table S1.

### Western blotting

Cell lysate in 1× SDS sample buffer was boiled for 5 min, separated by SDS-PAGE, and transferred to a PVDF membrane (Millipore) by electroblotting. After incubation with primary and HRP-conjugated secondary antibodies, the signals on the membranes were developed by a chemiluminescence reaction using the ImmunoStar Kit (Wako Chemicals), detected with a ChemiDoc imaging system (BioRad), and analyzed using ImageJ software (NIH). For analysis of the phosphorylation statuses of SR proteins, the samples were separated in a SDS-PAGE gel containing Phos-tag acrylamide (Fuji Film) and then transferred to a membrane according to the manufacturer’s manual. Antibodies are listed in Appendix Table S1.

### Quantitative and semi-quantitative RT-PCR

Total RNAs were prepared from cultured cells, subcellular fractions, or ChIRP samples using TRI Reagent or TRI Reagent LS (Molecular Research Center, Inc.), according to the manufacturer’s manual. The RNAs were treated with RQ1 RNase-free DNase (Promega) according to the manufacturer’s manual. First-strand cDNA was synthesized using High-Capacity cDNA Reverse Transcription Kits (Thermo Fisher Scientific). For quantitative reverse-transcription PCR (qRT-PCR), the sample and reference cDNAs were amplified using LightCycler 480 SYBR Green I Master Mix (Roche) and monitored using the LightCycler 480 System (Roche). For semi-quantitative RT-PCR, cDNAs were amplified by PCR to unsaturated levels, separated by electrophoresis, and stained with ethidium bromide. Images were obtained with a ChemiDoc system (BioRad) and analyzed with ImageJ software (NIH). Primers for PCR are listed in Appendix Table S2.

### Nascent RNA purification

Labeling and capturing of newly synthesized RNAs was performed using the Click-iT Nascent RNA Capture Kit (Thermo Fisher Scientific), according to the manufacturer’s protocol. Labeling was performed in the presence of 0.4 mM 5-Ethynyl Uridine (EU) for the indicated period. The purified RNAs were reverse-transcribed and analyzed by quantitative or semi-quantitative PCR as described above.

### FISH and immunofluorescence

For HSATIII RNA in situ hybridization, cells cultured on cover glasses were washed with ice-cold PBS and fixed in 4% PFA/PBS for 10 min at room temperature. Subsequently, the cover glasses were washed twice with ice-cold PBS and permeabilized in cold 70% ethanol for at least 1 h at 4°C. Next, the cover glasses were washed with 10% formamide/2× SSC, hybridized with a Dig-labeled HSATIII antisense oligonucleotide in hybridization buffer (10% formamide, 2× SSC, and 10% dextran) for 16 h at 37°C, incubated in 10% formamide/2× SSC for 30 min at 37°C, and then washed in 2× SSC for 5 min. After blocking in 3% BSA/TBST for 30 min, the cover glasses were incubated in 3% BSA/TBST containing an anti-dig antibody and an antibody against the protein of interest. After washing in TBST, the cover glasses were incubated in 3% BSA/TBST with Alexa 488- or Alexa 568-conjugated secondary antibodies. For immunostaining, the fixed cells were permeabilized in cold PBS containing 0.1% Triton X100 for 15 min, followed by the antibody reaction as described above. Images were obtained using a confocal laser scanning microscope, FLUOVIEW FV1000 (Olympus). Probes and antibodies are listed in Appendix Table S1.

### Bioinformatics analyses of RNA-seq data

Total RNAs of HeLa nuclear fractions were prepared as described above. Poly (A)+ RNA purification and library construction were performed using 200 ng of nuclear total RNAs and the Illumina TruSeq Stranded mRNA HT Sample Prep Kit. Subsequently, RNA-seq was performed using the Illumina Hiseq3000 system with the 36 bp single end method. Raw RNA-seq reads containing low quality and ambiguous bases were filtered out using cutadapt (version 1.9.1) (Martin, 2011) with the following parameters: -q 30 --max-n 0 -m 30. After this filtering, the remaining reads were mapped to the hg38 human reference genome using STAR aligner (version 2.5.3a) (Dobin, Davis et al., 2013) with the parameters --runThreadN 10 --outSAMtype BAM SortedByCoordinate --twopassMode Basic, and Gencode release27 annotation (Frankish, Diekhans et al., 2019).

For unambiguous counting of mapped reads with exon/intron resolution rather than transcript/gene levels, a single transcript was selected per gene using CGAT scripts (version 0.3.2) (Sims, Ilott et al., 2014) with the following parameters: gtf2gtf --method=filter --filter-method=representative-transcript. In these parameters, the “representative transcript” that shared the most exons with the other transcripts in the same gene was used. In the representative transcripts, the genomic coordinates of introns were calculated from the coordinates of exons using CGAT script with the parameters gtf2gtf --method=exons2introns, and these coordinates were merged into a single GTF file for the read counting process. For each intron annotation in the GTF file, a unique ID (ENSIxxxx; this ID is named after the upstream exon ID defined as ENSExxxx in the Gencode annotation) was added; “exon” was set into the feature field (third field in GTF format) for the counting process. The mapped reads were counted using the featureCounts algorithm (subread-1.4.6) (Liao, Smyth et al., 2014) with the following parameters: -s 2 -T 10 -t exon -g exon_id. Using these count data, differential expression analysis with exon/intron resolution was conducted using DESeq2 (version 1.10.1) (Love, Huber et al., 2014) with default parameters. Using the DESeq2 results, HSATIII target introns were selected according to the following criteria: fold-change (HSATIII-KD / control) <0.5 and adjusted *p*-value <0.01.

### Data analyses of RNA-seq and eCLIP

SRSF9 eCLIP peak data were obtained from the ENCODE website (ID: ENCFF432ASF, ENCFF327JJE) (Van Nostrand, Pratt et al., 2016). Regions that overlapped between the SRSF9 eCLIP peaks and the HSATIII target introns/adjacent exons were counted using bedtools (v2.28.0) (Quinlan & Hall, 2010) with the following parameters: intersect -s -c –F 0.5. As the control introns for the HSATIII target introns, overlaps between the SRSF9 peaks and all introns in the representative transcripts annotated in Gencode release 27 (for details, see above section “Bioinformatics analyses of RNA-seq data”) were also investigated.

Previously reported human (U-87 MG) or mouse (mESC) detained introns were obtained from the Supplemental Data of previous studies (Boutz et al., 2015; Braun et al., 2017). The genomic coordinates (based on hg19 or mm9) of these introns were converted to hg38 coordinates using the UCSC liftOver tool (Hinrichs, Karolchik et al., 2006) with default settings. The regions that overlapped between these introns and the HSATIII target introns were counted using bedtools with the following parameter: intersect –s.

### Gene ontology analysis

The gene ontology (GO) functional enrichment analyses were performed using DAVID Bioinformatics Resources 6.8 (LHRI).

### Quantification and Statistical analysis

For quantification, the experiments were performed as biological triplicate. Quantification of quantitative RT-PCR was performed using Lightcycler 480 software, version 1.5 (Roche). Quantification of semi-quantitative RT-PCR and western blot was performed using ImageJ software (NIH). The statistical significance of two-group and multiple comparisons was tested using GraphPad Prism7 software (GraphPad Software), as indicated in each figure legend. Non-adjusted (two-group comparison) and adjusted (multiple comparison) P-values were indicated in each figure.

## Acknowledgments

The authors thank Prof. Y. Suzuki (University of Tokyo) for conducting the RNA-seq analysis, Dr. N. Kataoka (University of Tokyo) for providing the SRSF9 expression plasmid, Dr. H. Maita (Hokkaido University) for providing the CHO cell line, and the members of the Hirose laboratory for valuable discussions. The computational analysis was partially performed on the NIG supercomputer at the ROIS National Institute of Genetics. This research was supported by MEXT KAKENHI Grant [to TH (JP26113002, JP16H06279, JP17H03630, and JP17K19335) and to NK (JP19K06478)] and the Tokyo Biochemical Research Foundation (to T.H.).

## Author contributions

KN and TH conceived and designed the study. KN conducted most of the experiments. SA and TN performed the mass spectrometry analyses. JI, GT, and KA analyzed the bioinformatics data from the RNA-seq and eCLIP experiments. KN and TH wrote the manuscript.

